# Optimizing Cognitive Phenotyping for Genetic Discovery in Alzheimer’s Disease: A Simulation-Based Evaluation of Mini-Mental State Examination-Based Decline Measures

**DOI:** 10.64898/2026.10.16.759741

**Authors:** Ruyu Shi, Eleanor Feingold, M. Ilyas Kamboh, Kang-Hsien Fan

## Abstract

**Background:** Alzheimer’s disease (AD) is a progressive neurodegenerative brain disorder characterized by cognitive decline. The Mini-Mental State Examination (MMSE) is a well-established screening tool that evaluates five cognitive domains with a total maximum score of 30. Although previous studies have suggested a genetic basis for cognitive decline in AD, inconsistent findings have been reported and may stem from variability in phenotypic definitions.

**Methods:** We conducted a simulation study using parameters informed by real-world AD cohorts to model 10-year trajectories of MMSE scores in a sample of 500 individuals. Three types of cognitive decline patterns—linear, quadratic, and sigmoid—were simulated. We incorporated realistic missing data mechanisms to reflect typical challenges in longitudinal data collection. From the simulated MMSE data, we derived seven distinct phenotypic measures of cognitive decline. Using statistical models that assumed additive genetic effects, we compared the power of these phenotypes across different missing data scenarios to identify the most effective strategies for phenotyping in genetic association studies.

**Results:** Adjusting for baseline MMSE improved the statistical power of measures that do not incorporate it in the definition. Across all simulated scenarios, simpler non-repeated measures, such as “Slope”, “dMMSEdt”, and “Time-to-MMSE10”, consistently outperformed the others.

**Conclusions:** The phenotypic measures “Slope” and “dMMSE/dt” consistently showed strong performance across a range of data distributions, while “Time-to-MMSE10” was particularly effective in cohorts with a higher proportion of individuals experiencing severe cognitive decline. These findings highlight the importance of tailoring the choice of cognitive decline measures to the specific characteristics of the dataset, such as the severity and distribution of decline, to improve the power to detect genetic factors associated with Alzheimer’s disease.

## Background

Alzheimer’s disease (AD), the most common form of dementia, is characterized by a chronic progressive decline of cognitive functions.^1^ The decline trajectories vary among individuals and are typically monitored using serial cognitive assessments. One of the most commonly used tools is the Mini-Mental Status Examination (MMSE), which assesses five key domains within approximately 15 mins. It yields an integer score ranging from 0 to 30, with lower scores indicating more severe cognitive impairment. In the advanced stage of the disease, typically reflected by an MMSE score below 9 or 10, the patient loses the ability to perform daily activities.^2,3^ Variations in scores below this threshold are generally considered to have limited clinical relevance. Because of this, the MMSE score is effectively limited to 21-22 clinically meaningful values. We refer to this type of measure as a semi-continuous phenotype: it is not categorical, as the score reflects the severity of cognitive function but lacks the granularity of a continuous variable. In addition to the MMSE, other neuropsychological battery tests that assess specific domains, such as memory, visuospatial ability, and executive function, often exhibit a similar semi-continuous nature.^4–7^ This feature makes it challenging to define an appropriate phenotype for measuring the cognitive decline rate in statistical analysis.

Genome-wide association studies (GWASs) have been widely used to identify genetic factors associated with AD. Since 2009, AD case-control GWASs have identified more than 95 risk loci ^8^. In contrast, research on the genetic basis of cognitive decline in AD cases has been more limited and has yielded inconsistent results mainly due to differences in how the cognitive decline phenotype is defined, measured, and analyzed across studies. For example, Wang et al. (2015) classified individuals as either slow or rapid progressors based on an arbitrary cutoff of a two-point annual decline in MMSE scores, and identified four potential risk loci (*PAX3, CCRN4L*, *PIGQ*, and *ADAM19*) at the suggestive significance ^9^. On the other hand, it can be treated continuously as a decline rate. However, inconsistent findings have emerged due to differences in how the decline rate is defined and the statistical methods used. Sherva et al. (2020) conducted a large GWAS involving 3,946 AD patients, using mixed-effect models and regressing raw longitudinal scores with a time interaction term to model the decline rate. Their analysis revealed several suggestive genetic associations on chromosomes 15 (*POLG, FAM214A*), 7 (*CNTNAP2*), and 3 (*lincRNA RP11-384F7*).^10^ Later, Homann et al. (2022) identified a single SNP within an intergenic region near *TNKS* on chromosome 8 that may influence the overall change of MMSE scores between the baseline and final visit.^11^

Prior research has emphasized the importance of phenotypic definition in identifying genetic associations. For example, a meta-analysis on Type-II diabetes showed that, beyond sample size, clearly defined and accurately measured phenotypic traits are essential for detecting susceptibility variants ^12^. In contrast, studies on AD cognitive decline differ widely in aspects such as the choice of neuropsychological tests, methods for calculating decline rates, follow-up durations, and statistical models. These variabilities make it difficult to compare results across studies and to identify the most reliable measures of cognitive decline.

Another major challenge in studying cognitive decline in patients is the presence of missing or censored data. Right censoring occurs when participants haven’t reached a cognitive milestone by the study’s end, often due to death or other complications. Left censoring occurs when participants join after the AD diagnosis point, resulting in the absence of early cognitive assessments. Additional issues, such as dropout, missed visits, or inability to complete tests, further complicate the analysis. These factors also complicate the development of a robust decline measure in longitudinal genetic studies. Therefore, evaluating different measures of cognitive decline is crucial to improve the statistical power of GWAS.

In this study, we conducted a simulation study by first extracting key parameters, such as age at initial visit, age of onset, baseline MMSE, and follow-up duration, from real AD cohorts. We then simulated 10-year MMSE trajectories for 500 individuals under linear, quadratic, and sigmoid decline patterns. To reflect realistic longitudinal data, we introduced various missing data mechanisms by masking simulated MMSE scores. Seven cognitive decline measures were then derived from the simulated MMSE scores. For each measure, we applied appropriate statistical models assuming additive genetic effects. By comparing the statistical power across phenotype measurements and missing patterns, we aim to identify and recommend the most effective calculation to represent cognitive decline phenotype for genetic association studies in AD. This work provides practical guidance for selecting outcome measures in future studies, with the goal of improving the detection of genetic risk factors and advancing our understanding of cognitive decline in AD.

## Methods

### Data Source and Key Parameter Estimation

Previous research has shown that individuals with AD exhibited diverse patterns and rates of cognitive decline.^13^ To capture this variability and simulate the full spectrum of MMSE trajectories, we used data frome two longitudinal AD cohorts: the University of Pittsburgh Alzheimer’s Disease Research Center (Pitt-ADRC, requested on Sep 1^st^, 2021) and the National Alzheimers’ Coordinating Center (NACC, Uniform Data Set (UDS) 2022 Freeze, requested on Sep 19^th^, 2022). Both cohorts have been previously described in detail.^13,14^

This study was an in-silico statistical simulation utilizing the parameters estimated from the previously collected, fully de-identified phenotypic data. No tissue samples or biospecimens were collected, nor were any new experiments conducted as part of this study. The original phenotype data were collected under protocols approved by the Institutional Review Boards (IRBs) from the original recruiting institutions, including the NACC and the Pitt-ADRC, and written informed consent was obtained from all participants or their legally authorized representatives. All data were fully de-identified before use and were assessed under appropriate Data Usage Agreements (DUAs) with NACC and Pitt-ADRC. All procedures complied with institutional and federal ethical guidelines.

The Pitt-ADRC cohort, which includes MMSE measurements following a clinical AD diagnosis, served as our primary data source. In contrast, the NACC cohort was used to supplement early-stage assessments near the time of AD onset (**Figure 1**).

**Figure 1.**
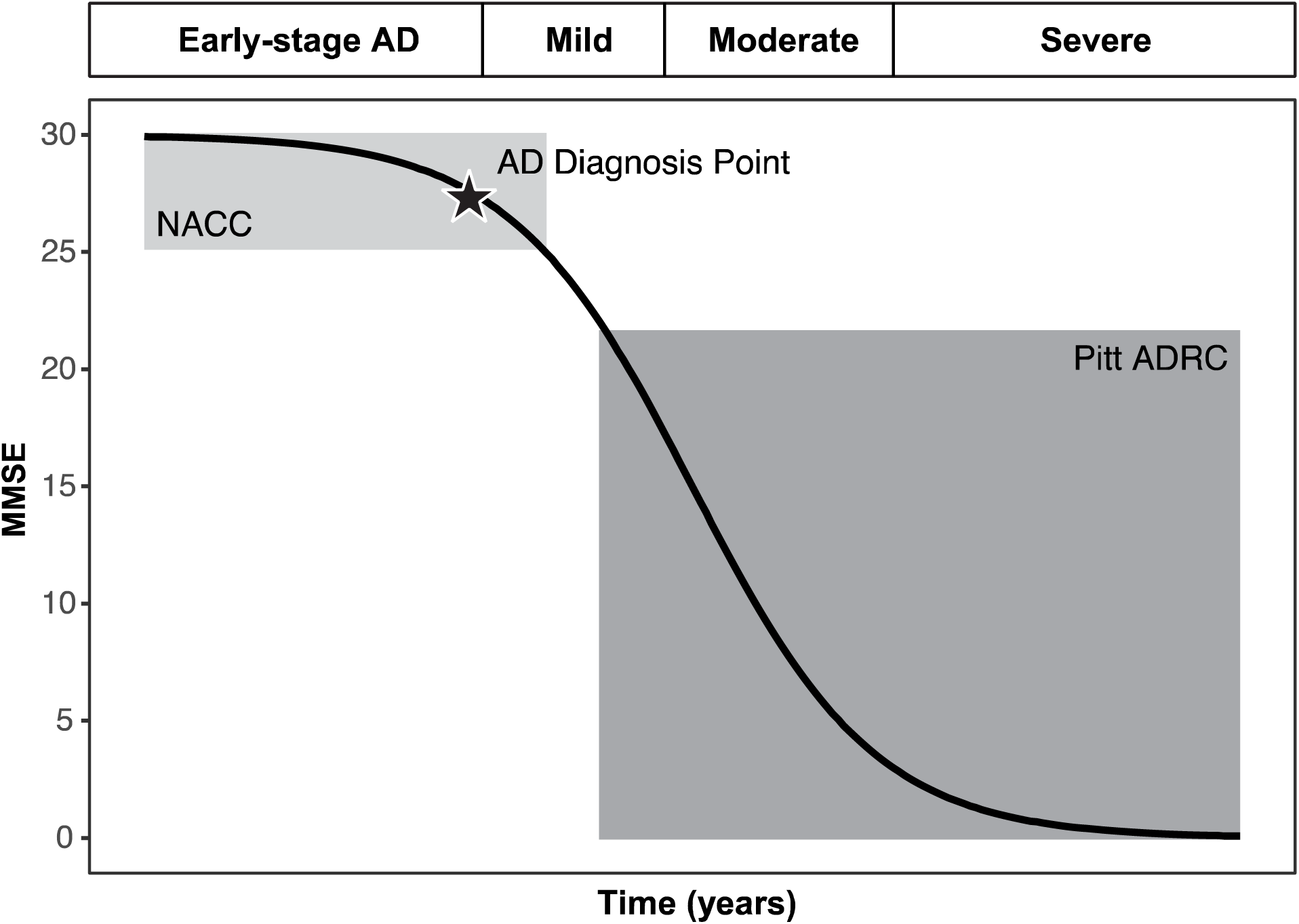
The illustration of hypothesized cognitive decline in the AD population

Demographic summaries of these two cohorts are presented in **Table 1**. The Pitt-ADRC included 1,739 individuals diagnosed with probable or possible AD, while 533 individuals from the NACC cohort who were cognitively normal at baseline and developed AD later were included to represent the transition phase.

**Table 1.**
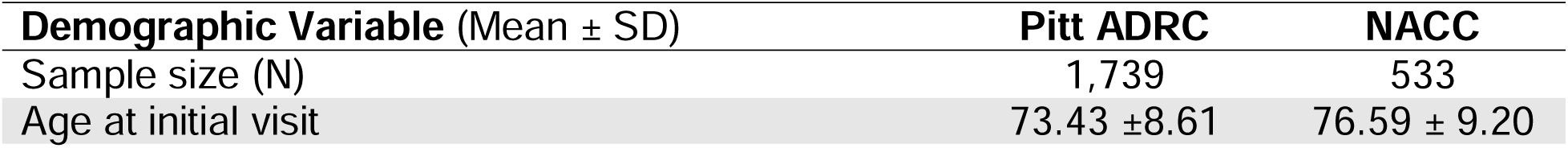
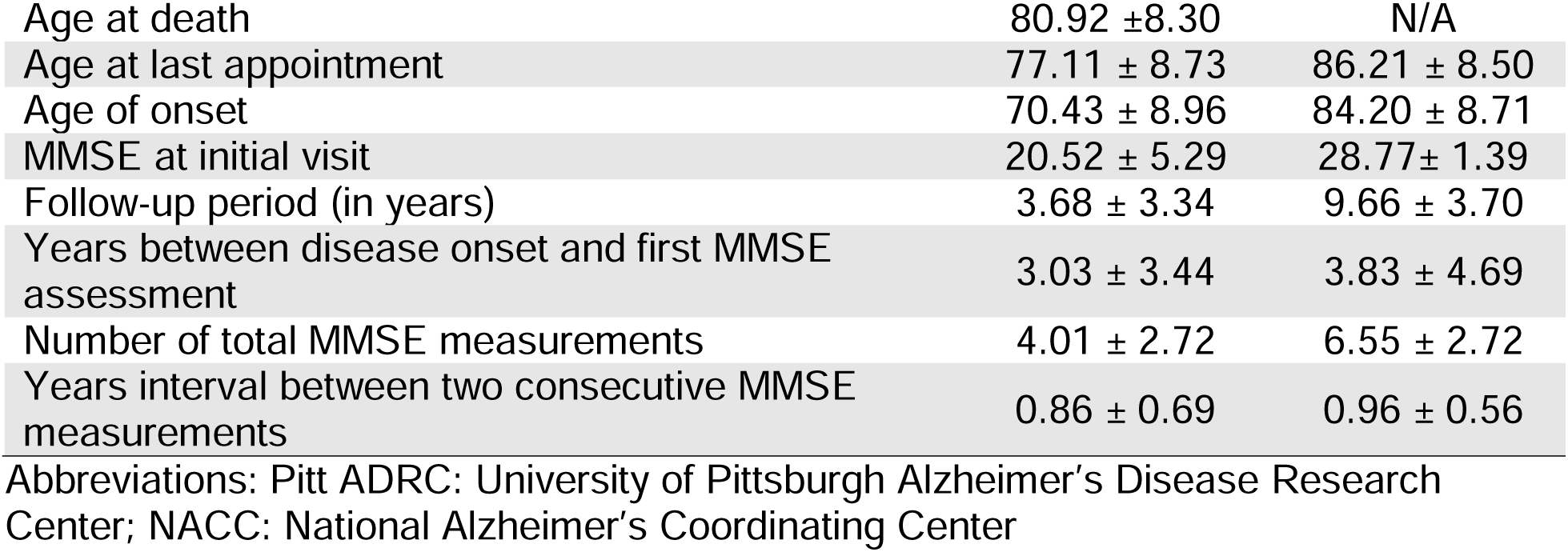
Demographics of the two source AD cohorts.

To combine the two cohorts in a reasonable way, we followed three main steps as illustrated in **Figure S1** [see Additional file 1]: (1) we applied linear mixed-effects model to MMSE scores from each cohort separately, with random effects for individuals and a fixed effect for time; (2) we identified the year when MMSE scores from both cohorts overlapped, and (3) retained MMSE scores from the NACC cohort before the overlap and from the Pitt-ADRC cohort after the overlap, aligning the Pitt-ADRC timeline accordingly. This process yielded a combined longitudinal MMSE dataset spanning a 10-year period, with a starting mean MMSE score of 28.71 (Standard Deviation, SD = 1.24).

Using the full combined dataset, we estimated parameters of cognitive decline under the assumption that MMSE scores decrease over time. Based on prior knowledge of AD cognitive trajectories, we modeled three distinct patterns of decline: linear, half-arch, and S-shaped, modeled using linear, quadratic, and sigmoidal functions, respectively (**Figure 2**). Linear and quadratic models were fit directly to the combined MMSE data to estimate trajectory parameters. To model the S-shaped trajectory, we employed a four-parameter logistic regression model, which enabled us to capture both the nonlinear curve and its linear decline phase. Details of each model and the key decline parameters are summarized in **Table 2**.

**Figure 2.**
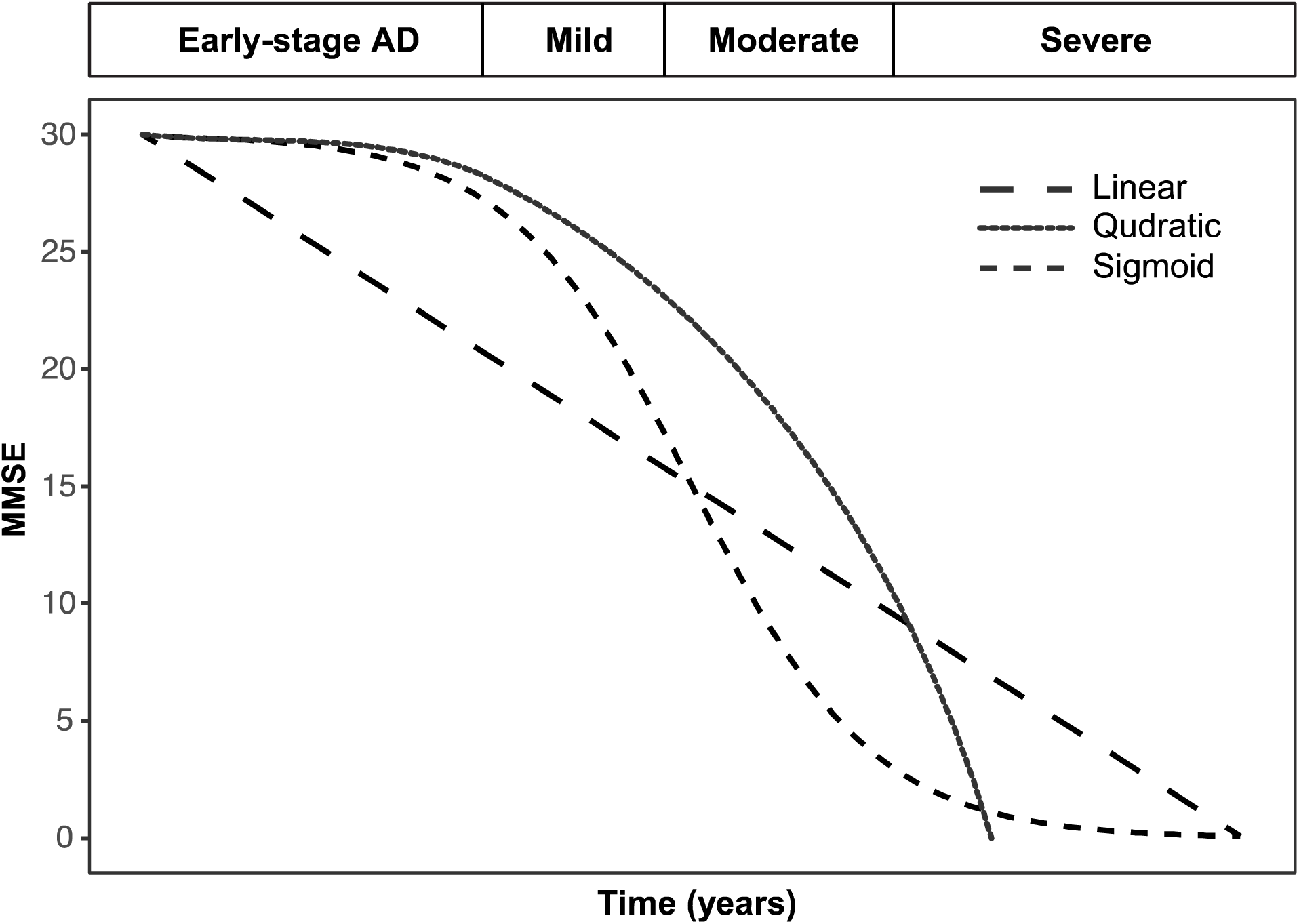
The illustration of three proposed cognitive decline trajectories over time

**Table 2.**
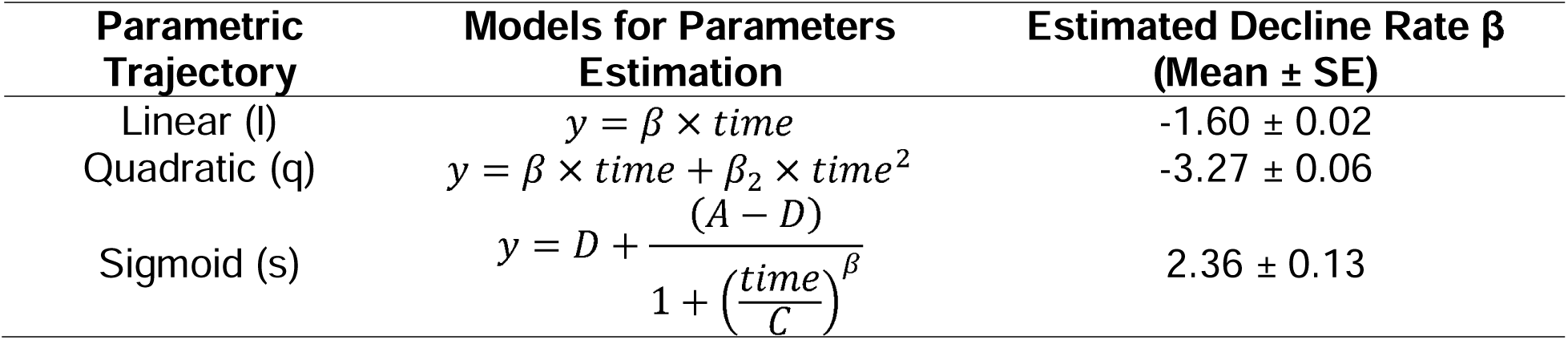
Parameters estimated from combined source AD cohorts.

| Parametric Trajectory | Models for Parameters Estimation | Estimated Decline Rate $\beta$ (Mean ± SE) |
| --- | --- | --- |
| Linear (l) | $y = \beta \times time$ | $-1.60 \pm 0.02$ |
| Quadratic (q) | $y = \beta \times time + \beta_2 \times time^2$ | $-3.27 \pm 0.06$ |
| Sigmoid (s) | $y = D + \frac{(A - D)}{1 + \left(\frac{time}{C}\right)^\beta}$ | $2.36 \pm 0.13$ |

### Simulation of MMSE Trajectories

Using the parameters in **Table 2** as reference, we tested multiple sets of decline rates under each proposed decline model, assuming an additive effect of genetic variant and a fixed sample size of 500. The relatively small sample size was chosen to better differentiate statistical power across models. Baseline MMSE scores were simulated from a truncated normal distribution with a mean of 28.71, a SD of 1.24, and an upper bound of 30. Genotypes were coded additively based on the number of minor (B) alleles, with values {0,1,2} representing genotypes AA, AB, and BB, respectively. The effect size of the susceptibility variant was defined as the regression coefficient associated with the number of minor alleles, assuming a linear trend per additional copy. Under this additive model, individuals with the AA genotype (no minor allele) were assigned the slowest rate of cognitive decline. Decline coefficients for the AB and BB genotypes were scaled proportionally to the number of minor alleles, with additional adjustments to the magnitude of these coefficients to facilitate statistical power comparisons across scenarios. Genotype group frequencies {AA, AB, BB} were determined based on a minor allele frequency (MAF) of 0.15, established through iterative simulations. To mimic the variability in assessment timing commonly observed in real-world longitudinal data, time intervals between MMSE assessments were randomly sampled within one standard deviation of the mean interval length from the source dataset. Each simulated individual has 10 MMSE assessments after this step. Additionally, to simulate the irregular timing of annual assessments observed in real-world data, the time intervals between consecutive MMSE assessments were randomly drawn within one SD of the mean interval length. We applied four different missing data mechanisms to the simulated datasets by masking simulated MMSE scores accordingly. First, we introduced a random missingness (RM) scenario by randomly masking MMSE measurements over time, reflecting unexpected missed visits. Second, we simulated left censoring (LCen) and right censoring (RCen) scenarios by randomly masking observations at the beginning or end of the trajectory, respectively. For LCen, we aimed to simulate scenarios where participants join after AD onset, resulting in missing early-stage cognitive measurements. Frequencies for masking were referenced on the time interval between onset and the first available MMSE assessment in the Pitt-ADRC cohort. For RCen, we simulated individuals with expected early dropout. Lastly, we simulated left and right censoring (LRCen) simultaneously to reflect more complex and realistic patterns of missing data. All the previously described settings were listed in **Table 3**.

**Table 3.** Details of simulated dataset scenarios.

| Experimental factor | Details |
| --- | --- |
| Sample size | 500 |
| Trajectory models<sup>a</sup> |  |
| Linear (l) | Decline coefficient 2.10, 2.15, 2.20 |
| Quadratic (q) | Decline coefficient 2.80, 2.83, 2.86 |
| Sigmoid (s) | Decline coefficient 3.00, 4.00, 5.00 |
| Missing data patterns under masking strategies: | Mask 1: Random missing (RM)<br>Mask 2: Left censoring (LCen)<br>Mask 3: Right censoring (RCen)<br>Mask 4: Left and right censoring (LRCen) |
| SNP minor allele frequency (MAF) | 0.15 |

<sup>a</sup> The decline coefficients of each parametric model are modified and finalized after comparing the power estimated from each cognitive measure, with the lowest value assigned to the genotype AA group, and the highest value assigned to the genotype BB group.

Finally, to reflect the clinical nature of MMSE scoring, all simulated scores were rounded to the nearest integer. This step ensured that the MMSE values for the 500 simulated individuals, each with varying trajectories and missing data patterns, closely resembled real-world clinical data. However, as previously noted, this rounding introduces what we refer to as a semi-continuous measurement, highlighting the need to evaluate the statistical power of various phenotype definitions and their corresponding analytical models. This motivates the analysis presented in the following section. We obtained the most realistic set of simulated MMSE scores at this stage, providing a strong foundation for downstream statistical analyses.

### Cognitive Decline Phenotype Construction and Statistical Analysis

We constructed seven distinct MMSE-based cognitive decline measures from the simulated longitudinal data, including 1) the regression slope of MMSE scores over time (“Slope”), 2) the total change between the first and last available MMSE score (“dMMSEdt”), 3) a binary classification of individuals as rapid or slow progressor based on a dMMSEdt threshold of 3 points (“Type”), 4) the full set of raw longitudinal MMSE scores, treated as repeated outcome (“MMSE”), 5) the absolute annual changes in MMSE scores between consecutive visits, treated as repeated outcome (“dMMSE_yr”), 6) the annual percentage changes in MMSE, treated as repeated outcome (“Perc_yr”), and 7) the time until an individual’s MMSE score drops to 10 or below, a milestone threshold often considered as lacking clinical interpretability as mentioned in the Introduction (“Time-to-MMSE10”) (**Figure 3**, **Table 4**). These phenotype definitions were either previously used in GWAS analyses or selected for their potential to serve as meaningful measures of cognitive decline when paired with appropriate statistical models. For the non-repeated measures, we apply linear regression to the “Slope” and “dMMSEdt” variables, logistic regression to the binary “Type” classification, and a Cox proportional hazards model to analyze the “Time-to-MMSE” variable. For the three longitudinal measures, “MMSE”, “dMMSE_yr”, and “Perc_yr”, we use generalized linear mixed-effect models.

**Figure 3.**
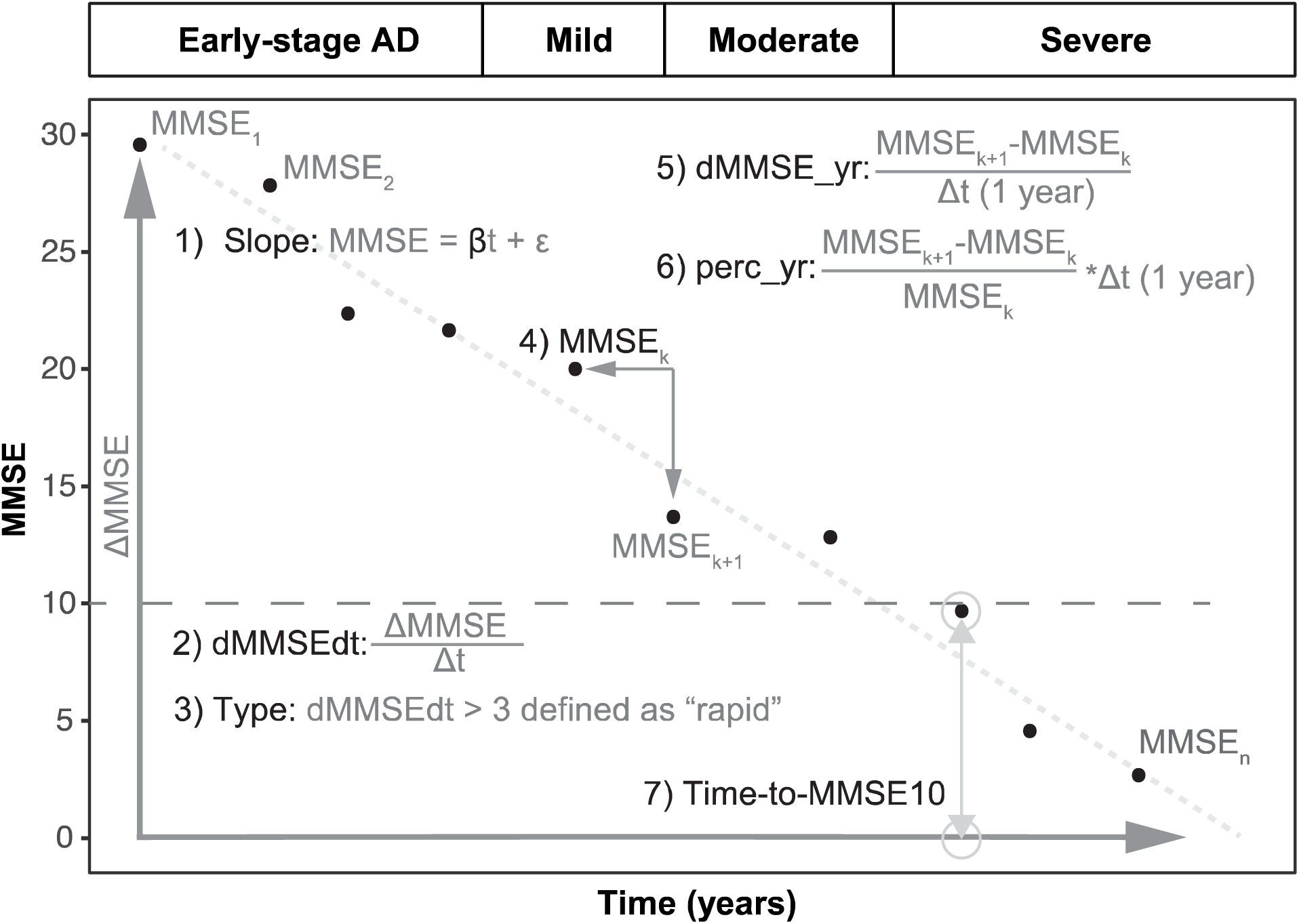
The illustration of seven MMSE-based cognitive decline phenotype measures calculation

**Table 4.**
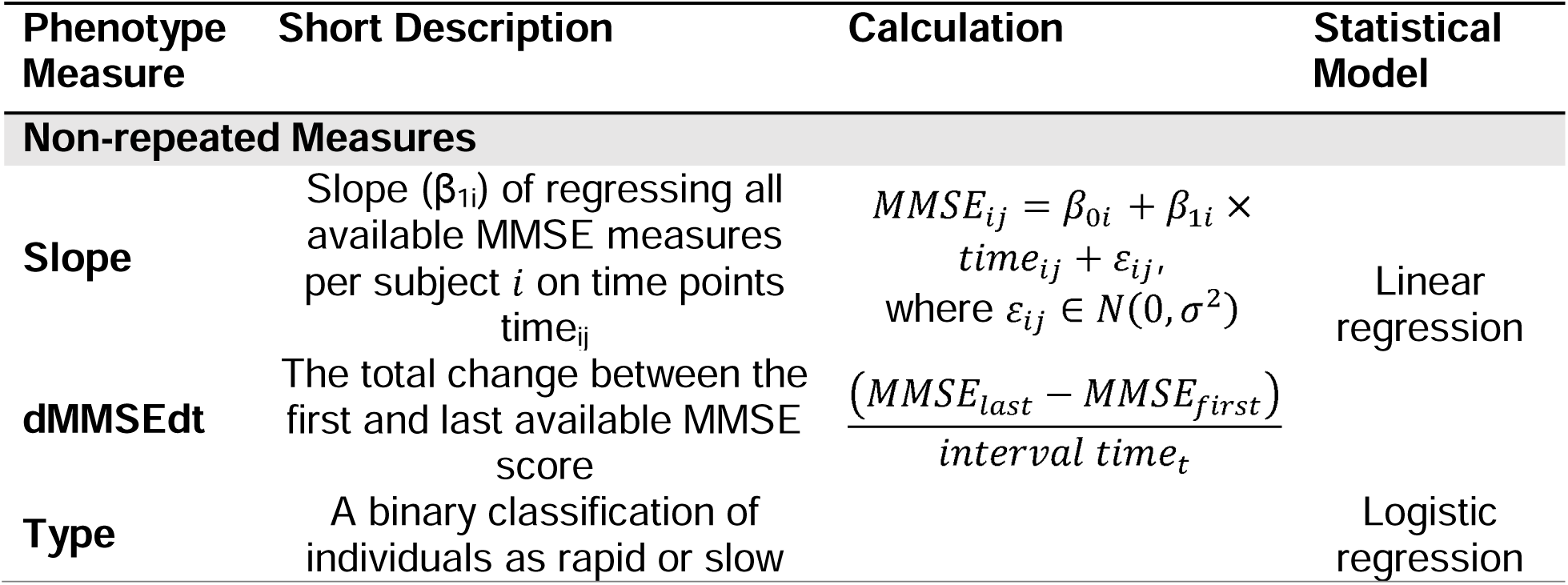
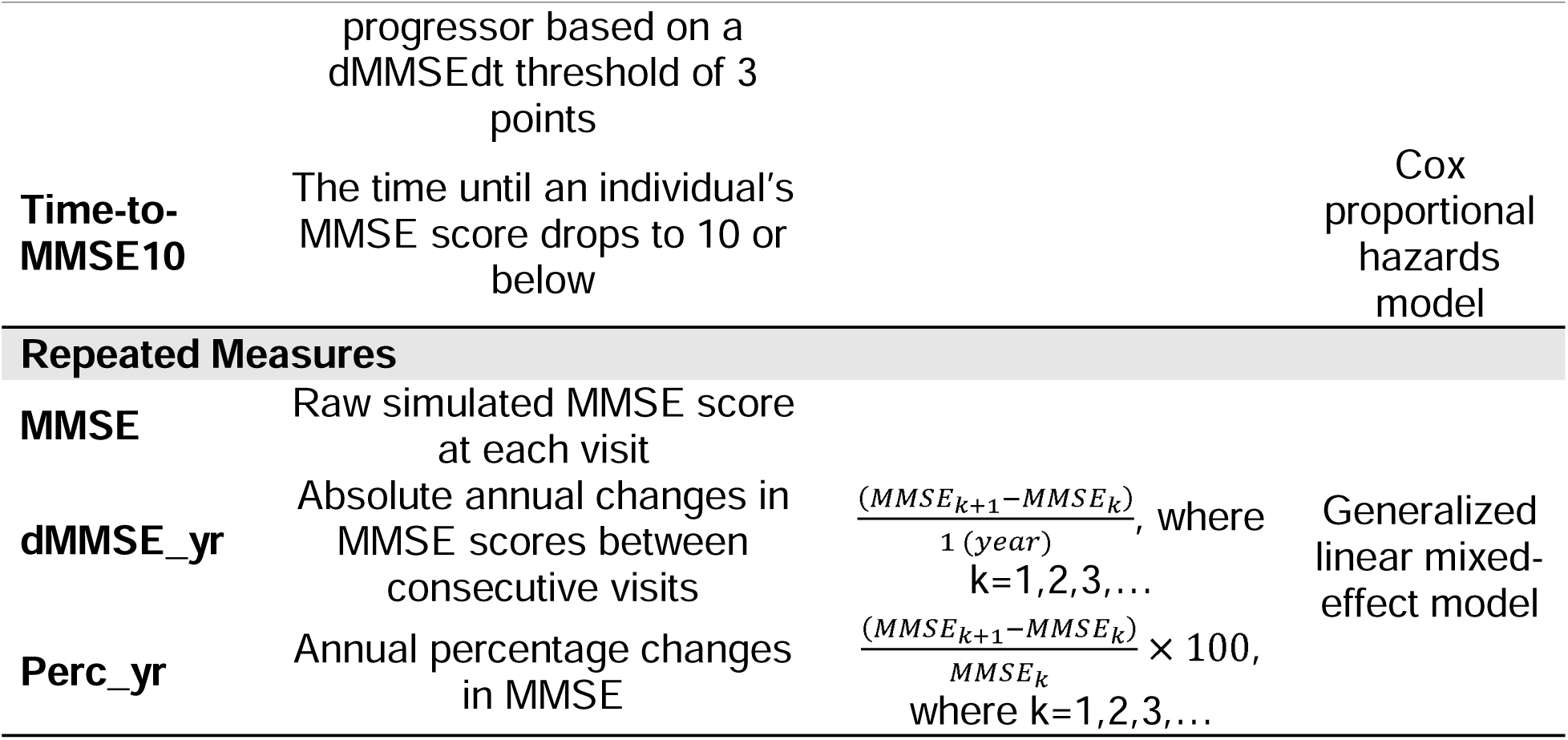
Cognitive decline phenotype definition and corresponding statistical model.

We performed 1,000 replicate analyses under each simulation scenario and calculated statistical power as the proportion of replicates under a p-value cutoff of 0.05. Analyses were conducted both with and without adjusting for baseline MMSE. The objective was to compare the performance of each MMSE-based cognitive decline measure and to identify measures that consistently demonstrate robust power across scenarios. All analyses were conducted using R Ver. 4.3.0.

## Results

**Figure S2** [see Additional file 2] illustrates the longitudinal MMSE trajectories under linear, quadratic, and sigmoid curves. **Table 5a** [see Additional file 3] summarizes descriptive statistics of simulated MMSE scores across different missing data scenarios. **Figure 4** and **Table 5b** [see Additional file 3] present the power comparisons of genetic association analyses for seven MMSE-based cognitive decline measures, covering all simulated scenarios that incorporate combinations of all trajectory types and missing data patterns.

**Figure 4.**
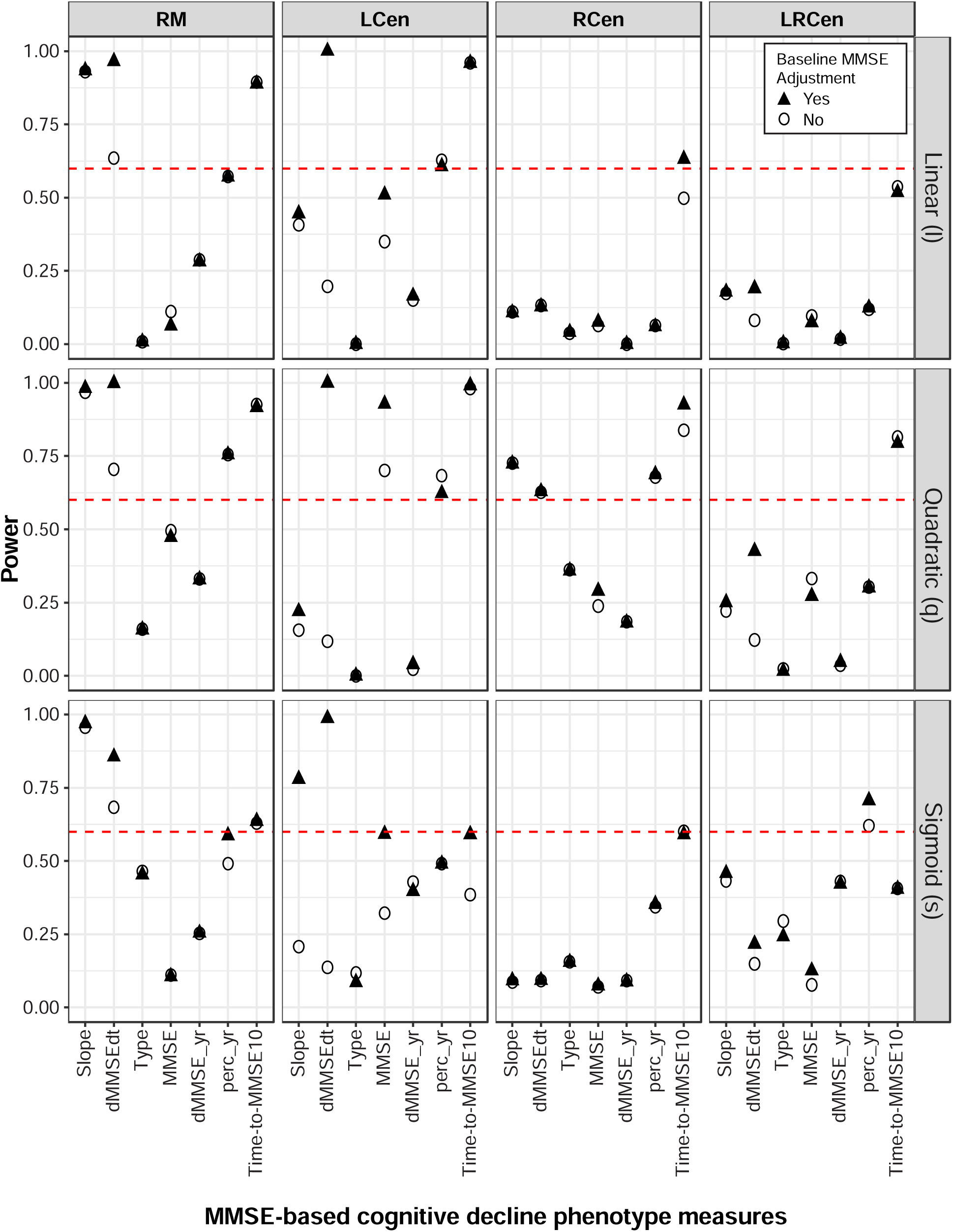
Statistical power comparisons of MMSE-based decline measures under various simulated data scenarios

Across all simulated scenarios, adjusting for baseline MMSE consistently improved the statistical power of every decline measure, especially for measures that did not initially incorporate it. However, the relative performance of decline measures varied depending on the specific simulation conditions. In general, when focusing on each individual block in **Figure 4**, where both the simulated trajectory and missing data pattern are held constant, the simpler, non-repeated measures, including “Slope”, “dMMSE/dt”, and “Time-to-MMSE10”, consistently outperformed the other four measures. This trend persisted within each missing data pattern, regardless of the trajectory (i.e., the three blocks in a column in **Figure 4**), where those simpler non-repeated measures continued to outperform others, including “Type”, “MMSE”, “dMMSE_yr”, and “Perc_yr” in terms of statistical power. When comparing across different missing data patterns under a fixed trajectory (i.e., the four blocks in a row in **Figure 4**), “Slope” and “dMMSE/dt” performed best in RM or LCen. However, in scenarios exhibiting extensive right censoring (RCen) or censoring from both ends (LRCen), the Cox regression model using the “Time-to-MMSE10” phenotype achieved the highest power.

Overall, these results suggest that simpler phenotype definitions and more straightforward statistical models tend to yield greater statistical power in detecting genetic associations related to cognitive decline. As expected, the binary “Type” measure consistently exhibited the weakest performance across all scenarios, likely due to the loss of information and reduced sensitivity inherent in dichotomizing continuous outcomes. Likewise, the three longitudinal repeated-measures approaches—“MMSE”, “dMMSE_yr”, and “Perc_yr”—consistently demonstrated lower power for different combinations of missing data patterns and trajectory types.

## Discussion

In this study, we began by estimating key parameters, such as the baseline MMSE score, using combined data from two longitudinal AD cohorts. Based on these estimates, we simulated MMSE-based cognitive trajectories under three distinct patterns of decline and incorporated various mechanisms for handling missing data. Using the rounded integer MMSE scores to mimic clinical assessments, we derived seven different phenotype measures and applied appropriate statistical models to assess their power. The rounding of MMSE scores, combined with their bounded scale (0–30) and the incorporation of diverse missing data patterns, resulted in semi-continuous measurements that may have obscured subtle nonlinear trends in cognitive decline, particularly evident in the quadratic trajectory blocks of **Figure S2** [see Additional file 2]. Nonetheless, we believe this artifact reflects the practical limitations commonly encountered in real-world studies using similar clinical data.

By mimicking key characteristics of MMSE data, this framework allowed us to comprehensively compare seven MMSE-based cognitive decline measures in AD populations, with an emphasis on statistical power relevant to downstream GWAS applications. We identified three phenotype measures—“Slope”, “dMMSEdt”, and “Time-to-MMSE10”—as having relatively high statistical power, even in small sample settings. Additionally, we found that adjusting for baseline MMSE scores significantly enhances the power of measures that do not inherently account for baseline performance. These findings align with statistical theory and echo conclusions from prior research on cognitive decline modeling.^15^

Among the three top-performing cognitive decline measures identified —“Slope”, “dMMSEdt”, and “Time-to-MMSE10”—the optimal choice depends on cohort characteristics known prior to analysis. These include the portion of the cognitive trajectory being studied, the length of follow-up, and the extent or nature of missing data. Based on our simulation results, “Slope” and “dMMSEdt” have consistently higher power in the LCen and RM scenarios. Their superior performance under these conditions suggests that both measures effectively utilize all available observations, regardless of follow-up duration. This makes them particularly well-suited for studies with irregular or incomplete data, increasing the likelihood of detecting meaningful cognitive changes despite data variability. “Time-to-MMSE10”, while ranking third in statistical power, addresses a distinct but complementary aspect of cognitive decline. Unlike “Slope” and “dMMSEdt”, which model continuous cognitive performance over time, “Time-to-MMSE10” focuses on the duration before reaching a milestone of severe impairment (MMSE ≤ 10). As such, it provides insight into the cumulative impact of decline and is especially valuable when the timing of critical cognitive deterioration is of interest. As shown in **Table 5b** [see Additional file 3], the statistical power of this measure increases with a higher proportion of individuals reaching the endpoint, as more events allow for clearer detection of effects influencing progression to severe impairment. Notably, this type of measure is also useful for studying transitions in cognitive functioning across disease stages, such as from cognitively normal to mild cognitive impairment (MCI) or from MCI to AD.^16,17^

In general, “Slope” and “dMMSEdt” offer robust performance across different data distributions, while “Time-to-MMSE10” is better suited for cohorts with a substantial proportion of severe cases. Between “Slope” and “dMMSEdt”, the two measures are equivalent when only two time points are available for a participant. However, “Slope” may yield more stable and robust estimates in datasets with many timepoints, as it leverages all available observations regardless of follow-up duration or specific timing. While this distinction was not critical in our motivating dataset, it may be important in other studies with more complex or extended longitudinal data.

Finally, we reiterate that all simulations and conclusions in this study were based on a design centered around the middle portion of the cognitive decline curve. However, our findings offer logical insights for other study types as well. For example, cohorts capturing early stages of cognitive decline—such as the transition from normal cognition to mild impairment—are likely to benefit from using “Slope” or “dMMSEdt”, but not “Time-to-MMSE10’, which is geared toward later-stage outcomes. In such early-stage studies, we recommend “Slope” in particular, as many participants may remain at the upper end of the MMSE scale for extended periods, reducing the sensitivity of dMMSEdt to capture subtle decline.

Conversely, in cohorts focused on normal aging with short follow-up durations, the MMSE may not be an ideal tool for tracking progression due to its limited sensitivity to detect mild cognitive changes. Prior studies suggest threshold MMSE scores of 24 for abnormal cognitive performance in dementia-free populations up to age 83, and 23.3 for individuals aged 90 and above.^18,19^

For studies that enroll participants later in the disease course, after significant decline, both “Slope” and “dMMSEdt” remain effective, with “dMMSEdt“ potentially offering an advantage due to the greater variability of MMSE scores at lower levels. Additionally, our conclusions can inform measure selection in studies using different cognitive scoring systems but targeting comparable phases of decline.

## Conclusions

In summary, to identify genetic variants associated with AD-related cognitive decline, we recommend using either non-repeated continuous measures (e.g., “Slope”, “dMMSEdt”) or time-to-event measures (e.g., “Time-to-MMSE10”), depending on the research question. Measures capturing the rate of decline emphasize linear trends over time, while time-to-event metrics highlight the cumulative progression to severe impairment. Importantly, the choice of decline measure should be informed by careful dataset screening. Key factors include the cognitive status of enrolled participants, baseline performance, and missing data patterns across the follow-up period. While our simulations were based on a relatively small dataset of 500 individuals—reflective of our discovery GWAS cohort with whole-genome sequencing—the conclusions are likely generalizable to larger cohorts with similar characteristics.

## Declarations

### Ethics approval and consent to participate

Not applicable

### Consent for publication

Not applicable

### Availability of data and materials

The data supporting the findings of this study are available from the National Alzheimer’s Coordinating Center (NACC) and the University of Pittsburgh Alzheimer’s Research Center; however, restrictions apply to the availability of these data, which were used under license for the current study and are therefore not publicly available. Data are, however, available from the authors upon reasonable request and with permission of the original site. The scripts and simulated datasets analysed during the current study are available from the corresponding author on reasonable request.

### Authors’ contributions

RS applied for the source data, conducted the data simulation and statistical analyses, generated the figures and tables, and contributed to drafting and revising the manuscript. EF and KF contributed to the study design, drafting of the manuscript, and critical revisions. MIK provided the source data and contributed to manuscript revisions. All authors read and approved the final version of the manuscript.

## Acknowledgements

The NACC database is funded by NIA/NIH Grant U24 AG072122. NACC data are contributed by the NIA-funded ADRCs: P30 AG062429 (PI James Brewer, MD, PhD), P30 AG066468 (PI Oscar Lopez, MD), P30 AG062421 (PI Bradley Hyman, MD, PhD), P30 AG066509 (PI Thomas Grabowski, MD), P30 AG066514 (PI Mary Sano, PhD), P30 AG066530 (PI Helena Chui, MD), P30 AG066507 (PI Marilyn Albert, PhD), P30 AG066444 (PI David Holtzman, MD), P30 AG066518 (PI Lisa Silbert, MD, MCR), P30 AG066512 (PI Thomas Wisniewski, MD), P30 AG066462 (PI Scott Small, MD), P30 AG072979 (PI David Wolk, MD), P30 AG072972 (PI Charles DeCarli, MD), P30 AG072976 (PI Andrew Saykin, PsyD), P30 AG072975 (PI Julie A. Schneider, MD, MS), P30 AG072978 (PI Ann McKee, MD), P30 AG072977 (PI Robert Vassar, PhD), P30 AG066519 (PI Frank LaFerla, PhD), P30 AG062677 (PI Ronald Petersen, MD, PhD), P30 AG079280 (PI Jessica Langbaum, PhD), P30 AG062422 (PI Gil Rabinovici, MD), P30 AG066511 (PI Allan Levey, MD, PhD), P30 AG072946 (PI Linda Van Eldik, PhD), P30 AG062715 (PI Sanjay Asthana, MD, FRCP), P30 AG072973 (PI Russell Swerdlow, MD), P30 AG066506 (PI Glenn Smith, PhD, ABPP), P30 AG066508 (PI Stephen Strittmatter, MD, PhD), P30 AG066515 (PI Victor Henderson, MD, MS), P30 AG072947 (PI Suzanne Craft, PhD), P30 AG072931 (PI Henry Paulson, MD, PhD), P30 AG066546 (PI Sudha Seshadri, MD), P30 AG086401 (PI Erik Roberson, MD, PhD), P30 AG086404 (PI Gary Rosenberg, MD), P20 AG068082 (PI Angela Jefferson, PhD), P30 AG072958 (PI Heather Whitson, MD), P30 AG072959 (PI James Leverenz, MD).

## List of abbreviations

AD: Alzheimer’s Disease
GWAS: Genome-Wide Association Study
LCen: Left Censoring
LRCen: Left and right censoring
MAF: Minor Allele Frequency
MCI: Mild Cognitive Impairment
MMSE: Mini-Mental State Examination
NACC: National Alzheimer’s Coordinating Center
Pitt-ADRC: University of Pittsburgh Alzheimer’s Disease Research Center
RCen: Right Censoring
RM: Random Missing
SD: Standard Deviation
UDS: Uniform Data Set

## Additional files

Additional File 1. Figure S1 Visualization of algorithms of combining two source AD cohorts

Additional File 2. Figure S2 Simulated longitudinal MMSE trajectories under linear, quadratic, and sigmoid models

Additional File 3. Table 5 Summary and Power Evaluation of Simulated MMSE Trajectories Across Scenarios

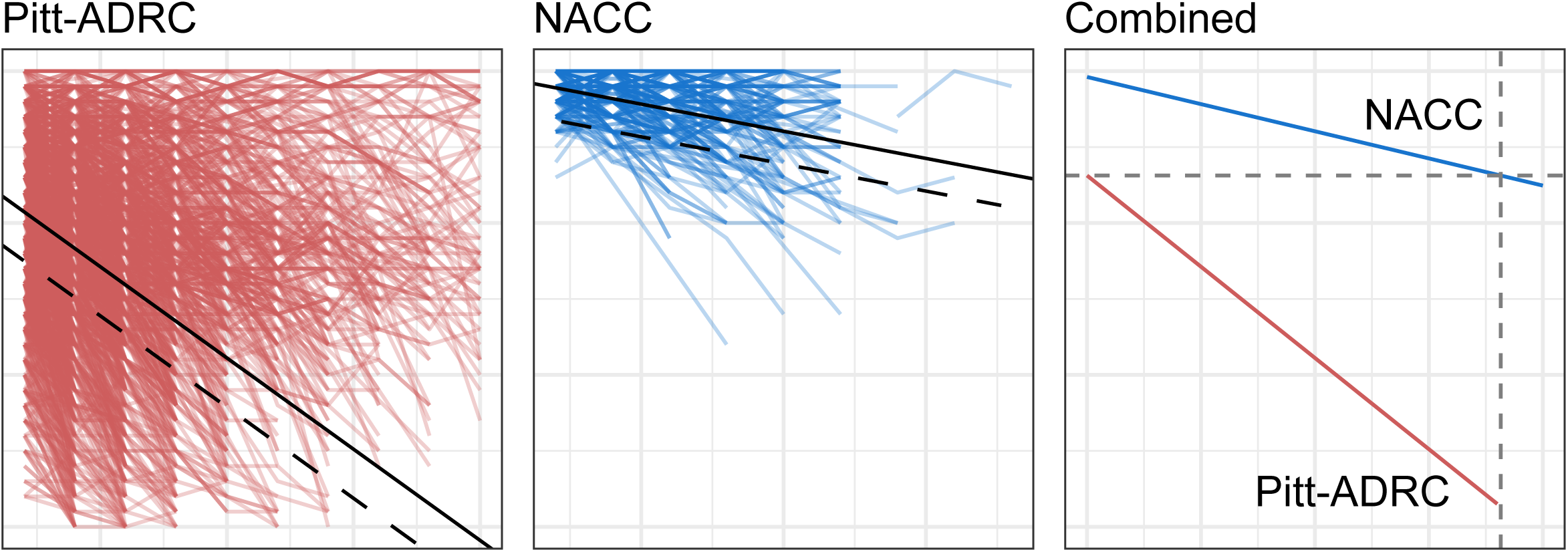

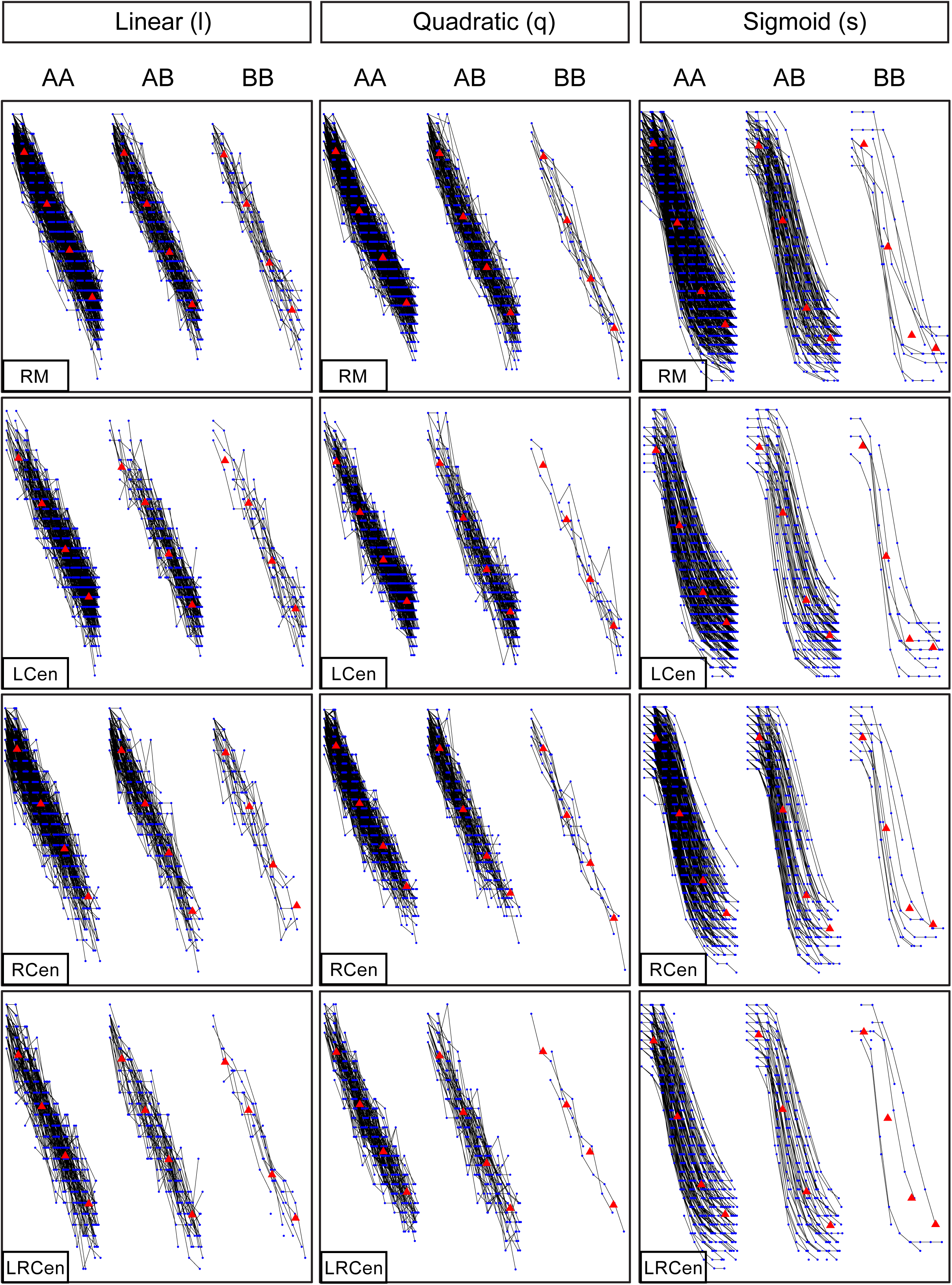

