## Supplementary material for "Optimizing Cognitive Phenotyping for Genetic Discovery in Alzheimer’s Disease: A Simulation-Based Evaluation of Mini-Mental State Examination-Based Decline Measures": Table 5

**Table 5. Summary and Power Evaluation of Simulated MMSE Trajectories Across Scenarios**

**Table 5a. Descriptive Statistics of Simulated MMSE Trajectories under each Scenario**

|  | Linear Line | | | Quadratic Curve | | | | | | | | | Sigmoid Curve | | | |
| --- | --- | --- | --- | --- | --- | --- | --- | --- | --- | --- | --- | --- | --- | --- | --- | --- |
|  | LRCen | Lcen | RM | | Rcen |  | LRCen | Lcen | RM | Rcen |  | LRCen | | Lcen | RM | Rcen |
| Baseline MMSE (Mean) | 23 | 18 | 27 | | 28 |  | 21 | 16 | 26 | 28 |  | 20 | | 15 | 26 | 27 |
| Last available MMSE (Mean) | 15 | 10 | 11 | | 18 |  | 13 | 8 | 9 | 16 |  | 10 | | 5 | 6 | 14 |
| Number of observations (Mean) | 5 | 5 | 6 | | 6 |  | 5 | 5 | 6 | 6 |  | 5 | | 5 | 6 | 6 |
| Proportion of Samples Reaching an MMSE of 10 or lower (%)^a^ | 19.2 | 68.6 | 47.1 | | 8.4 |  | 36.0 | 94.4 | 73.5 | 18.6 |  | 65.4 | | 98.7 | 91.3 | 44.6 |

**Table 5b. Power Comparisons under Simulated Scenarios after Adjusting for baseline MMSE^b^**

|  | Linear Line | | | Quadratic Curve | | | | | | | | | Sigmoid Curve | | | |
| --- | --- | --- | --- | --- | --- | --- | --- | --- | --- | --- | --- | --- | --- | --- | --- | --- |
|  | LRCen | Lcen | RM | | Rcen |  | LRCen | Lcen | RM | Rcen |  | LRCen | | Lcen | RM | Rcen |
| Slope | 17.8 | 44.6 | ***93.4*** | | 10.8 |  | 25.1 | 22.0 | ***98.2*** | 72.4 |  | 45.8 | | 78.0 | ***97.0*** | 9.2 |
| dMMSEdt | 19.0 | ***100.0*** | 96.6 | | 12.8 |  | 42.6 | ***100.0*** | 99.9 | 62.9 |  | 21.7 | | ***98.7*** | 85.6 | 9.3 |
| MMSE | 7.4 | ***51.0*** | 6.3 | | 7.6 |  | 27.2 | ***92.8*** | 47.3 | 29.0 |  | 12.6 | | ***59.2*** | 10.6 | 7.4 |
| dMMSE_yr | 1.8 | 16.4 | ***28.2*** | | 0.0 |  | 4.7 | 3.9 | ***32.9*** | 18.2 |  | ***42.2*** | | 39.7 | 25.4 | 8.9 |
| Perc_yr | 12.4 | ***60.7*** | 57.1 | | 6.0 |  | 30.1 | 62.3 | ***75.5*** | 68.8 |  | ***70.7*** | | 49.0 | 58.7 | 35.3 |
| Time-to-MMSE10 | 51.8 | ***95.9*** | 88.9 | | 63.2 |  | 79.4 | ***99.1*** | 91.7 | 92.6 |  | 40.5 | | 59.1 | ***63.7*** | 59.1 |
| Type | 0.2 | 0.0 | 0.8 | | ***4.1*** |  | 1.6 | 0.0 | 15.8 | ***35.9*** |  | 24.2 | | 8.6 | ***45.4*** | 15.5 |

Abbreviations:

1) LRCen – Left and Right Censoring; Lcen – Left censoring; RM – Random Missing; Rcen – Right censoring;

2) Slope – regression slope; dMMSEdt – overall MMSE differences calculated using the first and last MMSE divided by the follow-up years; dMMSE_yr – the MMSE differences per year; Perc_yr – the percentage change of MMSE per year; Time-to-MMSE10 – time until MMSE equals 10 is reached; Type – a binary slow or rapid outcome;

^a^The proportion listed here is the mean of the three genotype groups.

**^b^**The power estimates and proportions in this table are percentages, and the power estimates were adjusted for baseline MMSE.

The highest power of each MMSE-based measure under each trajectory is highlighted in ***bold and italic***.
