## Supplementary material for "Optimizing Cognitive Phenotyping for Genetic Discovery in Alzheimer’s Disease: A Simulation-Based Evaluation of Mini-Mental State Examination-Based Decline Measures": Table 4

**Table 4. Cognitive decline phenotype definition and corresponding statistical model**

| Phenotype Measure | Short Description | Calculation | Statistical Model |
| --- | --- | --- | --- |
| Non-repeated Measures | | | |
| Slope | Slope (β_1i_) of regressing all available MMSE measures per subject $i$ on time points time_ij_ | $MMSE_{ij}=\beta_{0i}+\beta_{1i}\times time_{ij}+\varepsilon_{ij}$,  where $\varepsilon_{ij}\in N\left( 0,\sigma^{2} \right)$ | Linear regression |
| dMMSEdt | The total change between the first and last available MMSE score | $\frac{\left( MMSE_{last}-MMSE_{first} \right)}{interval time_{t}}$ |  |
| Type | A binary classification of individuals as rapid or slow progressor based on a dMMSEdt threshold of 3 points |  | Logistic regression |
| Time-to-MMSE10 | The time until an individual’s MMSE score drops to 10 or below |  | Cox proportional hazards model |
| Repeated Measures | | | |
| MMSE | Raw simulated MMSE score at each visit |  | Generalized linear mixed-effect model |
| dMMSE_yr | Absolute annual changes in MMSE scores between consecutive visits | $\frac{\left( MMSE_{k+1}-MMSE_{k} \right)}{1 (year)}$, where k=1,2,3,… |  |
| Perc_yr | Annual percentage changes in MMSE | $\frac{\left( MMSE_{k+1}-MMSE_{k} \right)}{MMSE_{k}}\times100$, where k=1,2,3,… |  |
