## Supplementary material for "Optimizing Cognitive Phenotyping for Genetic Discovery in Alzheimer’s Disease: A Simulation-Based Evaluation of Mini-Mental State Examination-Based Decline Measures": Table 3

**Table 3. Experimental factors involved in simulated datasets**

| **Experimental factor** | **Details** |
| --- | --- |
| Sample size | 500 |
| Trajectory models^a^ |  |
| Parametric Linear (l) | Decline coefficient 2.10, 2.15, 2.20 |
| Parametric Quadratic (q) | Decline coefficient 2.8, 2.83, 2.86 |
| Parametric Sigmoid (s) | Decline coefficient 3,4,5 |
| Missing data patterns under masking strategies: | Mask 1: Random missing (RM) |
|  | Mask 2: Left censoring (LCen) |
|  | Mask 3: Right censoring (RCen) |
|  | Mask 4: Left and right censoring (LRCen) |
| SNP minor allele frequency (MAF) | 0.15 |

^a^Decline coefficients of each parametric model are modified and finalized after comparing the power estimated from each cognitive measure; with the lowest value assigned to the genotype AA group, and the highest value assigned to the genotype BB group.
