## Supplementary material for "Optimizing Cognitive Phenotyping for Genetic Discovery in Alzheimer’s Disease: A Simulation-Based Evaluation of Mini-Mental State Examination-Based Decline Measures": Table 2

**Table 2. Parameters estimation from source AD cohorts**

| **Parametric Trajectory** | **Models for Parameters Estimation** | **Estimated Decline Rate β**  **(Mean ± SE)** |
| --- | --- | --- |
| Linear (l) | $y=\beta\times time$ | -1.60 ± 0.02 |
| Quadratic (q) | $y=\beta\times time+\beta_{2}\times time^{2}$ | -3.27 ± 0.06 |
| Sigmoid (s) | $y=D+\frac{\left( A-D \right)}{1+\left( \frac{time}{C} \right)^{\beta}}$ | 2.36 ± 0.13 |
