## Supplementary material for "Optimizing Cognitive Phenotyping for Genetic Discovery in Alzheimer’s Disease: A Simulation-Based Evaluation of Mini-Mental State Examination-Based Decline Measures": Table 1

**Table 1. Characteristics of the two Motivating cohorts**

| **Demographic Variable** (Mean ± SD) | **Pitt ADRC** | **NACC** |
| --- | --- | --- |
| Sample size (N) | 1,739 | 533 |
| Age at initial visit | 73.43 ±8.61 | 76.59 ± 9.20 |
| Age at death | 80.92 ±8.30 | Not applicable |
| Age at last appointment | 77.11 ± 8.73 | 86.21 ± 8.50 |
| Age of onset | 70.43 ± 8.96 | 84.20 ± 8.71 |
| MMSE at initial visit | 20.52 ± 5.29 | 28.77± 1.39 |
| Follow-up period (in years) | 3.68 ± 3.34 | 9.66 ± 3.70 |
| Years between disease onset and first MMSE assessment | 3.03 ± 3.44 | 3.83 ± 4.69 |
| Number of total MMSE measurements | 4.01 ± 2.72 | 6.55 ± 2.72 |
| Years interval between two consecutive MMSE measurements | 0.86 ± 0.69 | 0.96 ± 0.56 |

Abbreviations: Pitt ADRC: –University of Pittsburgh Alzheimer’s Disease Research Center; NACC: –National Alzheimer’s Coordinating Center
